# Re-drawing Köppen-Geiger classes with microclimate: implications for nature and society

**DOI:** 10.1101/2023.11.20.567953

**Authors:** David H. Klinges, Ilya M. D. Maclean, Brett R. Scheffers

## Abstract

Scientists have long categorized the planet’s climate using the Köppen-Geiger (KG) classification to understand climate change impacts, biogeographical realms, agricultural suitability, and conservation. However, global KG maps primarily rely on macroclimate data collected by weather stations, which may not represent microclimatic conditions experienced by most life on Earth. Few studies have explored microclimate at broad scales, largely due to data and computational constraints. Here, we predicted KG classes separately from macroclimate and microclimate for over 32 million locations across six continents. Microclimate reclassified 38% of the total area, and microclimate KG classes were both more spatially variable, and encompassed a broader range of latitudes, relative to macroclimate KG classes. By redrawing the lines of climate classes, our study prompts a reevaluation of the importance of meteorological drivers of ecology across scales, shedding light on how natural, agricultural, and social systems experience and respond to global change.

## Introduction

Climate classification systems are a ubiquitous tool for understanding climate change and its impacts across space and time (Peel et al., 2007). The most prominent classification framework is the Köppen-Geiger (KG) system (Köppen, 1918; Geiger, 1954), which sorts the planet into major groups (A - E) subdivided into minor classes based upon annual averages and variability of temperature and precipitation (Figure 1). Global KG maps are widely used in education from primary school through graduate teaching, and inform research on crop suitability (Wang et al., 2022), species distributions (Mesgaran et al., 2014), human thermal comfort (Zhao et al., 2021) and disease spread (Savary et al., 2019).

**Figure 1.**
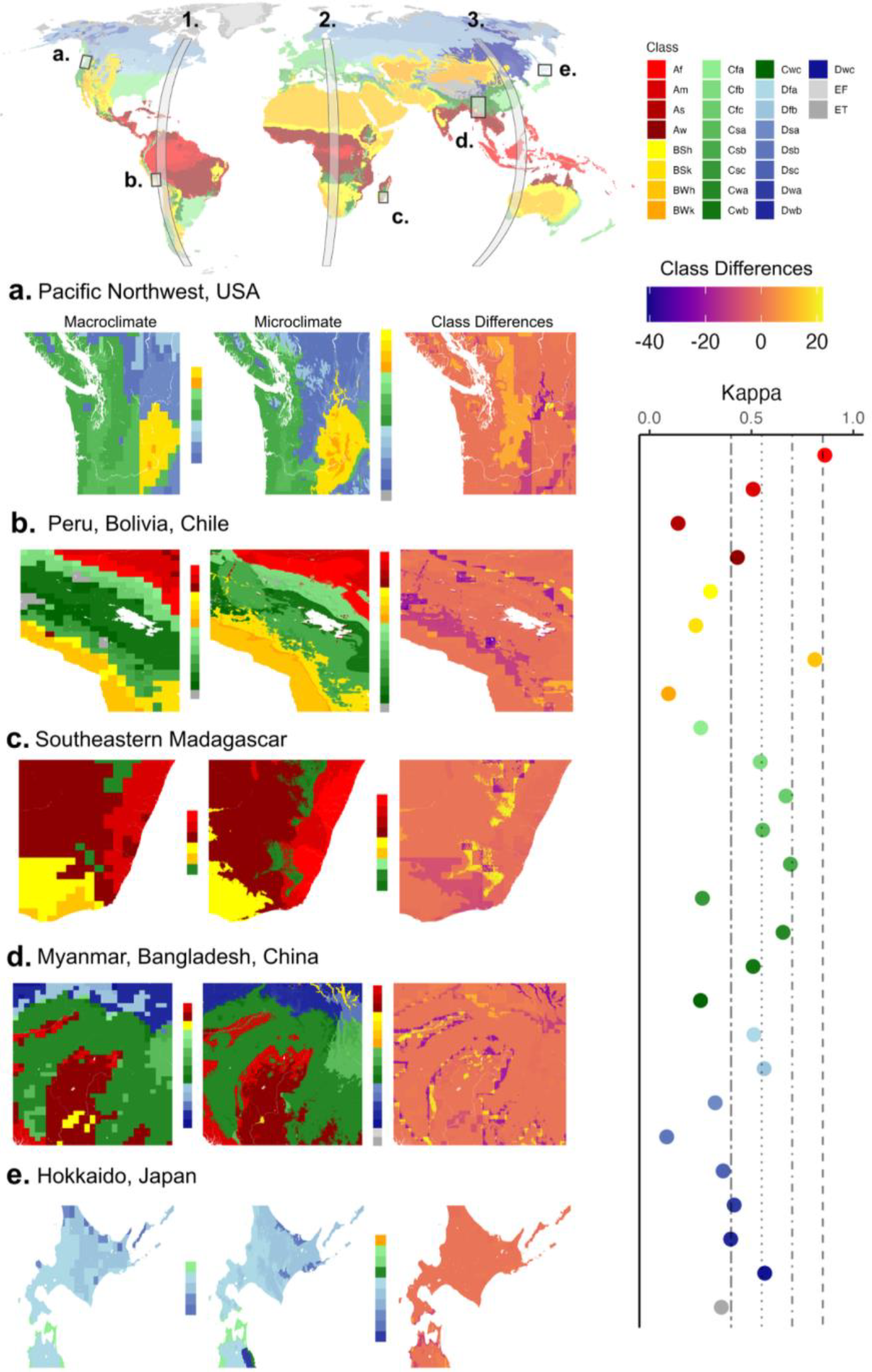
The Köppen-Geiger (KG) classification system subdivides the planet into discrete categories based upon annual averages and variability in temperature and precipitation. Top: Köppen-Geiger classes across the planet as represented by Kottek et al. (2006), displaying locations of case study regions (a - e) and latitudinal strips (1 - 3) for which we predicted KG classes from macroclimate and microclimate data. Left: maps of KG classes for each study region, with color legends depicting the sets of classes predicted by macroclimate and microclimate, and maps of the differences between macroclimate and microclimate class predictions. Right: amount of agreement, as measured by Fleiss’ kappa, between macroclimate and microclimate on the locations of Köppen-Geiger classes, with higher values indicating more agreement between macroclimate and microclimate. Dashed vertical lines indicate thresholds for (from left to right) poor, fair, good, very good, and excellent agreement; most classes (18 / 26) had poor or fair agreement, indicating large discrepancies between macroclimate and microclimate. Classes missing from this panel are those that were not predicted by macroclimate within case study regions or latitudinal strips.

Global KG maps (e.g. Kottek et al., 2006; Peel et al., 2007; Beck et al., 2018) have hitherto relied on macroclimate data obtained by weather stations situated 1.5 - 2 meters above the ground, away from topographic features, human developments, water, or vegetation (WMO, 2008). However, such climate data do not represent microclimates – the conditions shaped by local features (e.g. hills, valleys, and foliage) that impact near-surface heat and water exchange (Bramer et al., 2018). Microclimates are the broker of climate exposure for most terrestrial species, influencing physiology, community composition, and climate change-induced extinction risk (Suggitt et al., 2018). Given the discrepancies between regional macroclimate and local microclimate (De Frenne et al., 2019), macroclimate-derived KG classes may poorly represent the climatic reality for most ecosystems and human communities. Analyses using macroclimate KG maps have suggested that rice, corn, and millet croplands will need to shift considerably in space as large swathes of land become climatically unsuitable (Berg et al., 2013; FAO, 2021). Yet these efforts may be misguided as most crops are sensitive to fine-scale soil and air conditions– crops can be replanted into suitable microclimates a few meters away, rather than across hundreds of miles. Deriving KG classes from microclimate may also prominently reshuffle classes globally, revealing higher environmental heterogeneity (e.g. many KG classes expressed in a relatively small area) or anomalies relative to macroclimate (e.g. “tropical” KG classes located at high latitudes). Reclassifications based on microclimate could be immensely useful for better describing the climatic gradients experienced by plants and animals.

Here, we generated predictions of KG classes using macroclimate and near-surface microclimate, for a broad range of land uses and landforms globally. While an abundance of recent work has compared microclimate and macroclimate for particular ecosystems and taxa (Bramer et al., 2018), few have done so at the global extent. We discover that microclimatic KG classes dramatically diverge from macroclimate classes, exhibiting greater spatial variation and broader latitudinal ranges, which reshapes our understanding of the geography of climate. Re-evaluating climate classes, integrating how most life experiences climate, sheds light on the drawbacks of using macroclimate for understanding ecosystems and societies’ responses to global change.

## Methods

### Calculating Köppen-Geiger classes

We produced KG class maps, from macroclimate and microclimate data separately, using the classification system employed by (Beck et al., 2018) (WebTable 1). Macroclimate data, sourced from ERA5 (Hersbach et al., 2020), were hourly, 770-km^2^ datasets of precipitation and air temperature representative of conditions measured by free-standing weather stations. Microclimate data were 0.25-km^2^ resolution hourly predictions of 15-centimeter above ground air temperature and precipitation, subject to the influences of terrain and vegetation. Microclimate was derived using the microclimf microclimate model (Maclean, 2023) for temperature and interpolated CHELSA v2.1 data (Karger et al., 2021) for precipitation. The microclimf model is 1.58x more accurate than ERA5 at representing near-surface temperatures (Trew et al., 2023). We parameterized microclimf using the vegetation, terrain and climate forcing variables shown in WebTable 2. Given our focus on spatial rather than temporal comparisons, our maps represent conditions in 2015.

### Case Study Regions and Latitudinal Strips

We generated macroclimate and microclimate KG classes for five case study regions: the Pacific Northwest in the United States; areas of the Atacama Desert, Andean Altiplano, and Western Amazon basin (henceforth “Peru”, where most of this region falls); subtropical Southeastern Madagascar; South Asian lowlands and Himalayan foothills (henceforth “Myanmar”); and Hokkaido of Japan (Figure 1). To more thoroughly examine macro- and micro-KG classes across latitudes, we also generated KG classes for three discrete latitudinal strips from 60S° to 60N° in the Americas (75° W - 70° W), Africa and Europe (20° E - 22.5° E), and Oceania and Asia (115° E - 120° E).

### Analysis

We compared macroclimatic- and microclimatic-derived KG classes using three methods: cell-level class differences, latitudinal distributions and spatial variability. We measured the difference in KG classes, per spatial grid cell, when derived from macroclimate versus microclimate by using a simplified scoring system (Eccel et al., 2016): 10 for each change in major KG class (A to B = 10; A to C = 20), 1 for each first-degree subclass difference (Af to Am = 1), and 0.1 for each second-degree subclass difference (BSh to BSk = 0.1; WebTable 1). We calculated class differences both with microclimate at 0.25-km^2^ resolution and microclimate classes aggregated to their median and mode at 770-km^2^ resolution (to match macroclimate). For each class, we then calculated Fleiss’ Kappa for the level of agreement between micro- and macro-KG predictions (Fleiss, 1981).

Within each latitudinal strip, we compared the 2.5-97.5% percentile latitudinal range of each micro- and macro-KG class. We then identified poleward or equatorward extensions, defined as any location where a microclimate KG class lay outside the latitudinal range of its corresponding macroclimate KG class (demonstrated in Figure 2). For instance, if class Bf from macroclimate ranged between 12° and 45°, and class Bf from microclimate ranged between 7° and 53°, then the poleward extension was 53° - 45° = 8°, and the equatorward extension was 7° - 12° = -5°. We also tested for differences in the latitudinal distributions of each KG class using two-sample Wilcoxon tests.

**Figure 2.**
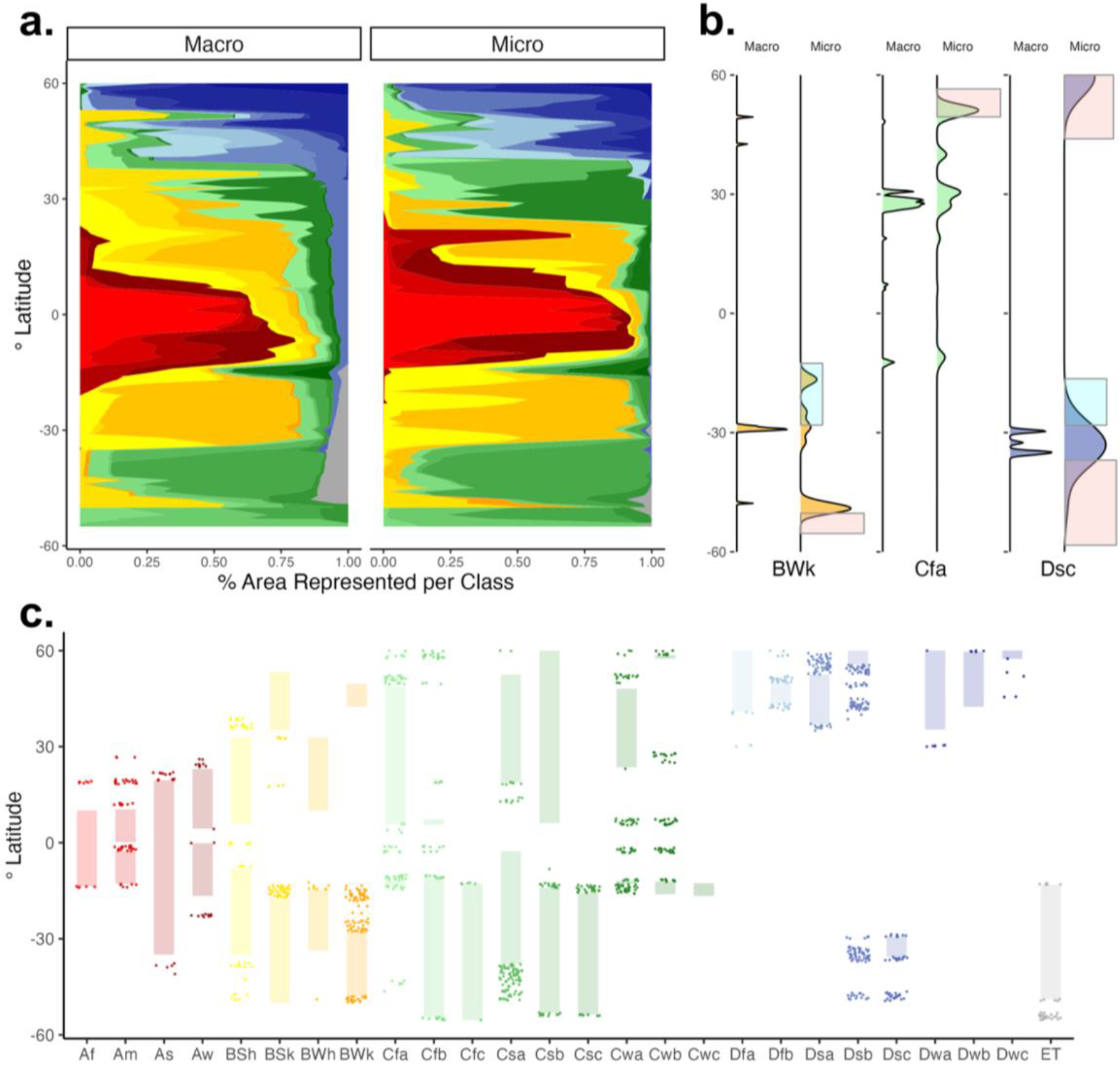
The latitudinal distributions of Köppen-Geiger classes from microclimate near the earth’s surface differ from the distributions of macroclimate-derived classes. (a) Density plots of each Köppen-Geiger class across latitude as derived from macroclimate and microclimate. (b) Density plots of three example classes (BWk, Cfa, Dsc) that demonstrate poleward extensions (shaded in pink) and equatorward extensions (shaded in light blue) of microclimate relative to macroclimate. (c) Microclimate extensions occurred for almost all classes. Here, lightly shaded bars are distributions of each macroclimate KG class in northern and southern hemispheres, with microclimate extensions represented as points above and below shaded bars (to reduce point density, each point represents all microclimate grid cells within 0.1° latitude of each other). Note that only the KG classes predicted in latitudinal strips by both macroclimate and microclimate are plotted in (c).

We measured spatial variability by calculating the number of macro- and microclimate KG classes falling within the same spatial area. To explore variability across spatial scales, we performed this calculation within sets of circles of increasing decimal degree diameter (0.005° to 16° for study regions, and from 1° to 180° for latitudinal strips; demonstrated in Figure 3). Across 50 randomly-placed sets of circles for each case study region and latitudinal strip, we calculated the mean number of macro- and micro-derived classes, and the 95% confidence intervals of each mean. We also repeated this analysis for all regions/latitudinal strips using microclimate classes averaged (median) to the same spatial resolution as macroclimate classes.

**Figure 3.**
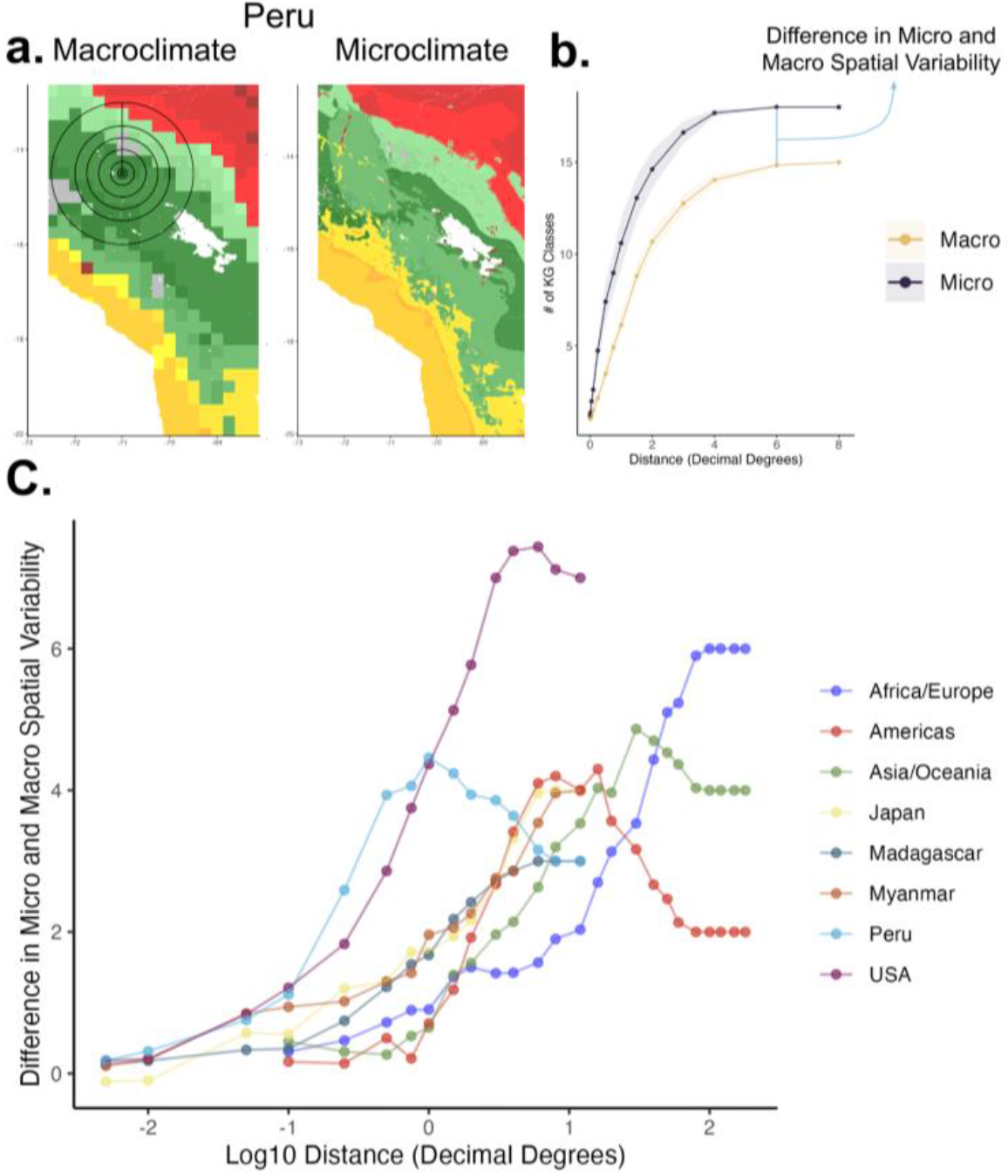
Predicting Köppen-Geiger classes with microclimate results in higher spatial variability in classes than predicting with macroclimate, and this trend holds across spatial scales. (a) For each case study region and latitudinal strip (here, using Peru as an example), we quantified the number of Köppen-Geiger classes within each of a set of increasing distances from a center coordinate (represented by concentric circles on the macroclimate panel, top left), separately for microclimate and macroclimate, and repeated this for 50 coordinates. (b) The mean number of classes for each distance (points), plotted with 95% confidence intervals of the mean (shaded ribbons surrounding points), demonstrates higher variability in microclimate classes than macroclimate classes across spatial scales. (c) The difference in spatial variability between microclimate and macroclimate classes for all study regions and latitudinal strips (here plotted across log_10_-transformed distances) was almost always positive, indicating consistently more variability in microclimate than macroclimate classes. This difference also increased with spatial area (increasing distance), peaking for most regions at 10 - 25 decimal degrees (∼ 1,000 - 2,000 km), suggesting that microclimate may play an important role for macro-scale ecological responses to climate.

## Results

Macroclimate- and microclimate-derived KG classes were considerably different (e.g., Figure 4). Across all five study regions and three latitudinal strips, 38% of cells differed by subclass (i.e. Af vs Am), 13% of cells differed by at least one major class (i.e. A vs. B), and 4% of cells differed by at least two major classes (i.e. A vs. C). D group classes (“continental”) on average diverged the most, with 52.6% of all macroclimate D cells designated as non-continental when derived by microclimate. Average agreement between macro-KG and micro-KG classes according to Fleiss’ Kappa was 0.435 (and 0.431 when the median microclimate class was taken for each macroclimate grid cell, WebFigure 1), which is considered “fair” but not “good” agreement (Fleiss, 1981). Agreement was highest for classes Af (0.86) and BWh (0.81), and lowest for classes Dsb (0.084), BWk (0.093), and As (0.14) (WebTable 3).

**Figure 4.**
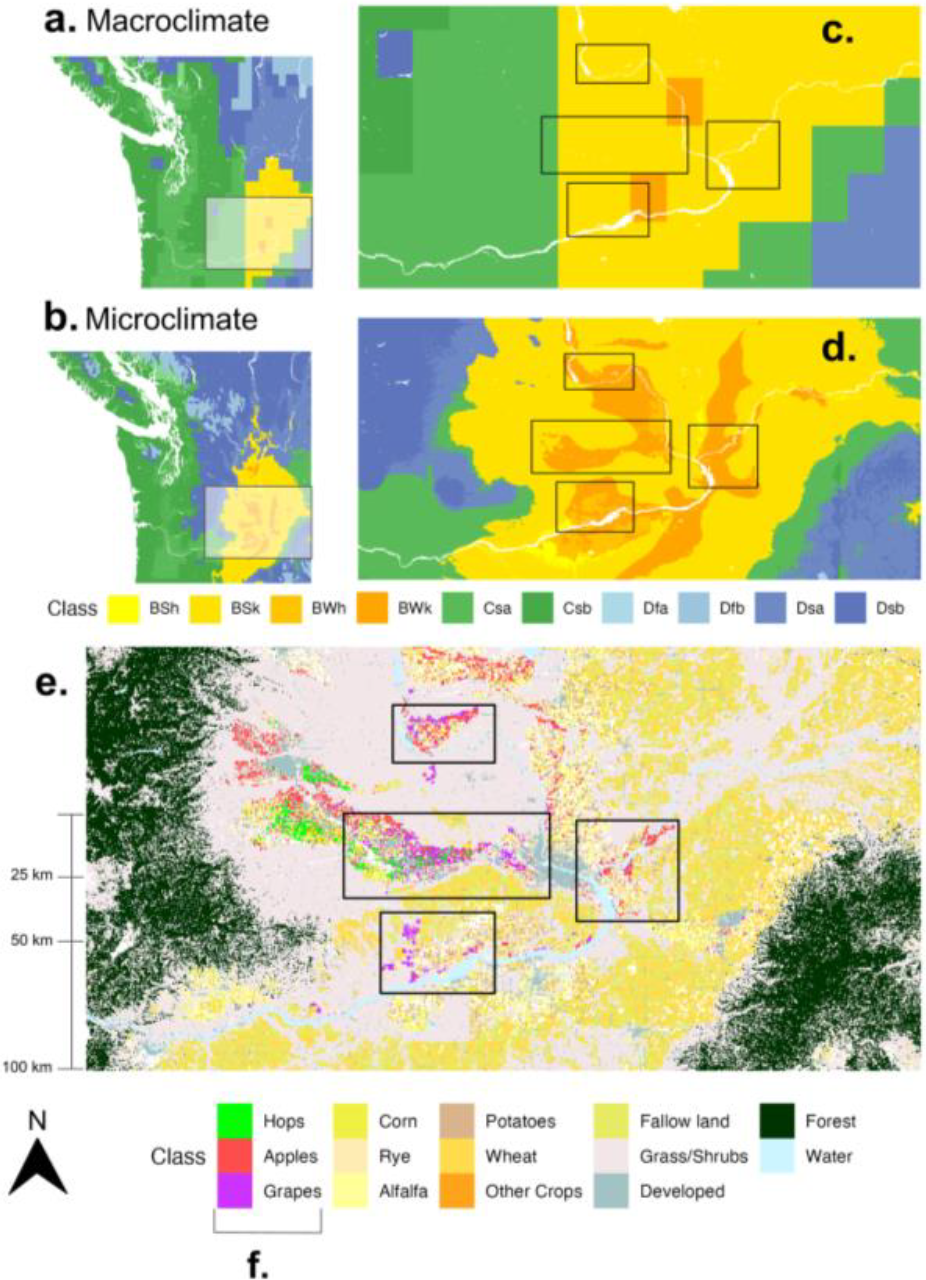
Köppen-Geiger classes as predicted by microclimate reflected gradients of land use better than classes predicted by macroclimate. In the Pacific Northwest of the USA, climate regimes at large scales are driven by distance to coasts, elevation, and latitude, which are generally reflected in class predictions by both macroclimate (a) and microclimate (b). However as shown in insets from Eastern Washington state (c-d), microclimate’s higher spatial resolution and better proximity at representing near-surface conditions generate class discrepancies. In the left side of (d), microclimate predictions reflect how the Cascades Mountain Range cause D-group classes (“continental” classes) to infiltrate to lower latitudes, while the coarser-resolution elevation model underlying macroclimate data (c) did not reflect the range’s topographic complexity, thereby predicting only a more subtle change from Csa to Csb (two classes that only differ based upon summer temperatures, not precipitation regimes). Within the Columbia River Basin (center of c and d scenes), one of the most commercially important agricultural regions in the USA, microclimate Köppen-Geiger classes more closely reflect fine-scale changes in agricultural land use (e, USDA-NASS, 2023). Microclimate predicts swathes of BWk (“cold desert climate”) and BWh (“hot desert climate”), which are preferred climate zones for planting regional strains of wine grapes, hops for brewing beer, and apples (f), given that desert-like conditions help keep disease-causing fungi and other pathogens at bay. Black boxes in (c - e) indicate areas of cash crop production in BWk and BWh climates. Microclimate successfully predicted BWk and BWh climates for 23.8% of the land where these three crops are planted, while only 3.9% of such cropland was predicted as BWk by macroclimate (and macroclimate failed to predict BWh at all). Not all lands of suitable microclimate are planted with cash crops, in part due to constraints from residential development and irrigation. Given that the Pacific Northwest accounts for 99% of all USA hops acreage (and 25% of global acreage) amounting to $662 million in revenue in 2021, the accuracy and scale of climate class maps used for interpreting present and future climate suitability can have significant economic ramifications.

The distribution of latitudes of each class was on average 2.94° broader, and centered 2.3° farther from the equator, when derived from microclimate than macroclimate. On average for each class within latitudinal strips, 9,046 km^2^ of microclimate predictions were extensions, i.e. fell outside the range of macroclimate predictions for the same class. Poleward extensions were more frequent, and of higher magnitude, than equatorward extensions (Figure 2). Wilcoxon tests indicated significant differences in latitudes between macroclimate and microclimate for all classes, with effect sizes ranging from very small (0.01, class Dwb) to moderate (0.33, class As; WebTable 3).

Microclimate KG classes also consistently demonstrated higher spatial variability than macroclimate KG classes for all study regions (Figure 3), even when microclimate was aggregated to the same spatial resolution as macroclimate (WebFigure 2). Notably, the difference in spatial variability between micro- and macro-classes increased with spatial scale (Figure 3c).

## Discussion

### Persistence disagreement between macroclimate and microclimate Köppen-Geiger classes

Globally pervasive differences in the composition and configuration of Köppen-Geiger classes were evident when derived from near-surface microclimate rather than free-air macroclimate, even when both were at the same spatial resolution. Given that most life on earth experiences microclimates instead of ambient macroclimate, these discrepancies entail meaningful errors in knowledge drawn from prior macroclimate classifications. Transitioning to the next generation of microclimate KG classes could serve biological research, conservation, and land planning.

The disagreement between macro- and micro-derived KG classes manifests in real differences in land use and agricultural suitability, as is clear in the Pacific Northwest of the United States (Figure 4). Here, the regional cash crops of apples, wine grapes, and beer hops are specifically planted in cool desert climate (BWk) and hot desert climate (BWh) to lower disease pressure (Smith, 2001). Microclimate successfully predicted BWk and BWh climates for 23.8% of the land where these three crops are planted, while only 3.9% of such cropland was predicted as BWk or BWh by macroclimate. When predicting future land suitability for such climate-sensitive crops, relying on macroclimate may lead to inaccurate predictions (e.g. minimal future area of suitable regional climate despite many suitable microclimates) and risk losses in the billions USD (Houston et al., 2018).

### Microclimate expands latitudinal distributions of Köppen-Geiger Classes

We found that 86% (24/28) of KG classes had broader latitudinal distributions when derived using microclimate (Figure 2). Of these 24 classes, an average of 11.2% of the total surface area of each microclimate class fell outside the latitudinal range of that same class delineated using macroclimate. If this extension of KG-classes across latitude remained constant across the terrestrial planet, then the global area of latitudinal extensions per KG class would be 232,555 km^2^, or roughly the area of the state of Oregon (USA) or the country of Ghana. Broader latitudinal distributions of micro-KG classes also suggested the need for some class rebranding. For instance, microclimates of class Cfa, or “humid subtropical”, were abundant above 50° latitude, far beyond latitudes typically considered “subtropical”.

For 64% of all classes, the equatorward latitudinal range limit of micro-KG classes extended further than the corresponding macroclimate KG class. These equatorward extensions generally entail cooler microclimates than the regional macroclimate and may thus provide refugia against contemporary climate change (Dobrowski, 2011). Poleward extensions were even more prevalent (71% of all KG classes), and encompassed larger areas (Figure 2). While these warmer extensions situated amidst cooler regional climate may afford opportunities for certain agricultural crops, they may hinder the dispersal for wild species tracking their thermal niches as the climate warms (Senior et al., 2019). Therefore such microclimates found beyond macroclimate latitudinal ranges may either facilitate, or disrupt, species’ responses to climate change.

### Spatially variable microclimate across scales

Micro-classes had consistently higher spatial variability than macro-classes across scales (Figure 3), and this held even when microclimate was coarsened to match the resolution of macroclimate (WebFigure 2). For our microclimate classes, the average distance from any cell to the nearest different major class was 4.14 km, while for macroclimate this average distance was 127.12 km. Climate classification systems have often been employed to group broad areas of the globe, yet when microclimate is considered, small extents of a few square kilometers may contain multiple classes. For example the city of Cusco (Peru) represented six microclimate classes ranging from tropical monsoon to mediterranean warm/cool summer climates (Am, As, Bsh, Cfa, Csa, Csb); macroclimate predicted only one class here. Upgrading climate classifications by using microclimate may enhance city planning to reduce human thermal mortality, given different risks faced by those near urban green spaces than in urban heat sinks (Aram et al., 2019).

With such fine spatial variability in climate, many mobile organisms could easily traverse multiple microclimatic KG classes in a single day as a thermoregulatory response to climatic extremes (Woods et al., 2015). In addition, individuals of the same population may persist in separate KG classes, leading to local physiological adaptations. This also helps explain why the directions of species’ range shifts often do not follow macroclimate gradients across elevation and latitude (Maclean & Early, 2023) or why climate-sensitive species are less prone to extirpation in topographically heterogeneous locations (Suggitt et al., 2018).

Micro-derived KG classes were also more spatially variable than macroclimate across continents. Across distances of 10,000 km, microclimates on average expressed four more KG classes than macroclimate (Figure 3). This suggests microclimate is important for even macroecology. For instance, the difference in climate variability experienced by tropical versus temperate organisms, situated thousands of kilometers apart, may have more to do with what microclimates they occupy (e.g. forest understories versus open grasslands) rather than the change in macroclimate across latitude (Klinges and Scheffers 2021). We encourage further exploration of how microclimate shapes physiology, behavior, and evolution across broad scales (Kearney, 2020).

### Improving the Utility of Climate Classification Systems

Our study, in conjunction with growing evidence of the importance of biometeorology to ecology, suggests that macroclimate-derived KG classes do not accurately reflect the composition, nor the configuration, of climate regimes as experienced by most life on earth. These differences were not just due to spatial resolution, as macroclimate classes did not match microclimate classes even when at the same spatial resolution. Furthermore, it is well-known that microclimate varies vertically as well as horizontally. Climate classifications may therefore be more useful if developed in three dimensions rather than two, or at least if using climate measurements from the height(s) relevant to the target species and processes.

Updated approaches of classifying climate that capture ecologically-relevant variation can help inform conservation and forecast the effects of climate change. Spatial variation in microclimate at both small and large scales points to its relevance when estimating climate connectivity, or informing restoration to improve connectivity (Senior et al., 2019). Predicted changes in climate classes have been used to understand rates of climate change (Kottek et al., 2006) and climate impacts on future agricultural suitability (Wang et al., 2022), carbon stocks (Gibson et al., 2021), and human thermal comfort (Mishra & Ramgopal, 2013)– yet all of such work has relied on macroclimate. Using microclimate is paramount for establishing relevance of forecasts to most ecosystems and species, especially as microclimates may warm faster or slower than ambient macroclimate (Maclean et al., 2017; De Lombaerde et al., 2022). Furthermore, new “ultra-tropical” or “ultra-desert” classes may need to be defined as the world experiences non-analog conditions (Trew et al., 2023). Although we urge caution when using climate classes derived from macroclimate, climate classification systems in general are still useful heuristics for simplifying multiple climate variables (e.g. temperature and precipitation) into single categories, and communicating where and when do consequential changes in climate occur (i.e. a shift from one class to another).

## Conclusion

We have shown that when compared to macroclimate-derived Köppen-Geiger classes, microclimate classes are consistently different, have higher spatial variability across scales, and not all “tropical” and “polar” classes are found just in the tropics or near the poles, thereby redrawing the lines of climate classes across the planet. Macroclimate remains adequate for understanding the climatic “backdrop” of a region, but researchers and practitioners must remain cautious in assuming that all species within a given macroclimate class experience similar climate regimes. Most climate-relevant management decisions, such as planning cities or planting crops, are also best informed by microclimate data. The influence of microclimates on ecosystems and species distributions is not ephemeral nor only felt at local scales, and may significantly impact macroecology. By recognizing the unique patterns and broad consequences of microclimates, we can refine our understanding of multiscale climate-driven ecological dynamics.

## Acknowledgements

We thank J. A. Baecher, L. Evans and L. Soifer for helpful conversations and insights. D.H.K. was supported by the National Science Foundation Graduate Research Fellowship (DGE-1842473). I.M.D.M. was supported by the Natural Environment Research Council (NE/L00268X/1). B.R.S. was supported by an Alfred P. Sloan Fellowship.

## Open Research statement

The following two options both apply to this manuscript:

Data are already published and publicly available, with those items properly cited in this submission: ERA5 data are available for download from:

https://cds.climate.copernicus.eu/cdsapp#!/dataset/reanalysis-era5-single-levels?tab=form

The microclimf microclimate model is publicly available in a GitHub repository:

https://github.com/ilyamaclean/microclimf

Data are provided for peer review (shared either privately or publicly in a repository).

Raw data and code for reproducing analyses and display items are available in the following Zenodo repository:

https://zenodo.org/record/8372921

