## Supplemental Files for "Re-drawing Köppen-Geiger classes with microclimate: implications for nature and society"

WebTable 1. Köppen-Geiger classification schematic used here, reflecting the modified version used by Beck et al. (2018). Classes are divided into 1st (major), 2nd, and 3rd degree subdivisions, each represented by a numeric value (Eccel *et al.* 2016), which were used to quantify differences in classes between macroclimate and microclimate. Also provided are the description for each class and the criteria by which the class is calculated. Variable definitions: MAT = mean annual air temperature (°C); Tcold = the air temperature of the coldest month (°C); Thot = the air temperature of the warmest month (°C); Tmon10 = the number of months with air temperature >10 °C (unitless); MAP = mean annual precipitation (mm y^−1^); Pdry = precipitation in the driest month (mm month^−1^); Psdry = precipitation in the driest month in summer (mm month^−1^); Pwdry = precipitation in the driest month in winter (mm month^−1^); Pswet = precipitation in the wettest month in summer (mm month^−1^); Pwwet = precipitation in the wettest month in winter (mm month^−1^); Pthreshold = 2 × MAT if >70% of precipitation falls in winter, Pthreshold = 2 × MAT+28 if >70% of precipitation falls in summer, otherwise Pthreshold = 2 × MAT+14. Summer (winter) is the six-month period that is warmer (colder) between April-September and October-March.

| **1st** | **2nd** | **3rd** | **Value** | **Description** | **Criteria** |
| --- | --- | --- | --- | --- | --- |
| A |  | | | Tropical | Not (B) & *Tcold*≥18 |
|  | f |  | 11 | Rainforest | *Pdry*≥60 |
|  | m |  | 12 | Monsoon | Not (Af) & *Pdry*≥100-*MAP*/25 |
|  | s |  | 13 | Savannah dry summer | Not (Af) & *Pdry<*100-*MAP*/25 & P*dry* in summer months |
|  | w |  | 14 | Savannah dry winter | Not (Af) & *Pdry<*100-*MAP*/25 & P*dry* in winter months |
| B |  | | | Arid | *MAP*<10×*Pthreshold* |
|  | S | h | 21.1 | Hot Steppe | *MAP*≥5×*Pthreshold & MAT≥18* |
|  | S | k | 21.2 | Cold Steppe | *MAP*≥5×*Pthreshold & MAT<18* |
|  | W | h | 22.1 | Hot Desert | *MAP*<5×*Pthreshold & MAT≥18* |
|  | W | k | 22.2 | Cold Desert | *MAP*<5×*Pthreshold & MAT<18* |
| C |  | | | Temperate/Mesothermal | Not (B) & *Thot*>10 & 0<*Tcold*<18 |
|  | f | a | 31.1 | Humid subtropical | Not (Cs) or (Cw) & Thot≥22 |
|  | f | b | 31.2 | Oceanic | Not (Cs) or (Cw) or (a) & Tmon10≥4 |
|  | f | c | 31.3 | Subpolar oceanic | Not (Cs) or (Cw) or (a) or (b) & 1≤Tmon10<4 |
|  | s | a | 32.1 | Mediterranean hot summer | *Psdry<40* & *Psdry<Pwwet/3 & Thot≥22* |
|  | s | b | 32.2 | Mediterranean warm/cool summer | *Psdry<40* & *Psdry<Pwwet/3 & not (a) & Tmon10≥4* |
|  | s | c | 32.3 | Mediterranean cold summer | *Psdry<40* & *Psdry<Pwwet/3 & not (a) or (b) & 1≤Tmon10<4* |
|  | w | a | 33.1 | Dry-winter humid subtropical | *P* *wdry<P* *swet/10 & Thot≥22* |
|  | w | b | 33.2 | Dry-winter subtropical highalnd | *P* *wdry<P* *swet/10 & not (a) & Tmon10≥4* |
|  | w | c | 33.3 | Dry-winter cold subtropical highalnd | *P* *wdry<P* *swet/10 & not (a) or (b) & 1≤Tmon10<4* |
| D |  | | | Cold | Not (B) & *Thot*>10 & *Tcold≤*0 |
|  | f | a | 41.1 | No-dry season hot summer continental | Not (Ds) or (Dw) & Thot≥22 |
|  | f | b | 41.2 | No-dry season warm summer continental | Not (Ds) or (Dw) & not (a) & Tmon10≥4 |
|  | f | c | 41.3 | No-dry season subarctic or boreal | Not (Ds) or (Dw) & not (a, b, or d) |
|  | f | d | 41.4 | No-dry season subarctic or boreal with severe winter | Not (Ds) or (Dw) & not (a or b) & Tcold<-38 |
|  | s | a | 42.1 | Dry hot summer continental | *Psdry<*40 & *Psdry<Pwwet*/3 *& Thot≥22* |
|  | s | b | 42.2 | Dry warm summer continental | *Psdry<*40 & *Psdry<Pwwet*/3 *& not (a) & Tmon10≥4* |
|  | s | c | 42.3 | Dry summer subarctic or boreal | *Psdry<*40 & *Psdry<Pwwet*/3 *& not (a, b, or d)* |
|  | s | d | 42.4 | Dry summer subarctic or boreal with severe winter | *Psdry<*40 & *Psdry<Pwwet*/3 *& not (a or b) & Tcold<-38* |
|  | w | a | 43.1 | Dry winter hot summer continental | *Pwdry<Pswet*/10 *& Thot≥22* |
|  | w | b | 43.2 | Dry winter warm summer continental | *Pwdry<Pswet*/10 *& not (a) & Tmon10≥4* |
|  | w | c | 43.3 | Dry winter subarctic or boreal | *Pwdry<Pswet*/10 *& not (a, b, or d)* |
|  | w | d | 43.4 | Dry winter subarctic or boreal with severe winter | *Pwdry<Pswet*/10 *& not (a or b) & Tcold<-38* |
| E |  | | | Polar | Not (B) & *Thot*≤10 |
|  | T |  | 51 | Tundra | *Thot*>0 |
|  | F |  | 52 | Frost/Ice cap | *Thot*≤0 |

#### WebReferences

Beck, H.E., Zimmermann, N.E., McVicar, T.R., Vergopolan, N., Berg, A. & Wood, E.F. (2018) Present and future Köppen-Geiger climate classification maps at 1-km resolution. *Scientific Data*, **5**, 180214.

Eccel E, Zollo AL, Mercogliano P, and Zorer R. 2016. Simulations of quantitative shift in bio-climatic indices in the viticultural areas of Trentino (Italian Alps) by an open source R package. *Computers and Electronics in Agriculture* **127**: 92–100.

WebTable 2. Input data products, and the variables extracted from them, used to parameterize the *microclimf* microclimate model.

| **Product** | **Variable(s) used** | **Spatial Resolution** | **Spatial Extent** | **Temporal Resolution** | **Temporal Extent** | **Citation** |
| --- | --- | --- | --- | --- | --- | --- |
| ECMWF ERA5 reanalysis | 2m temperature, 2m dewpoint temperature, surface pressure, 10m u wind, 10m v wind, total precipitation, total cloud cover, mean surface net longwave radiation flux, mean surface downward longwave radiation flux, total sky direct solar radiation at surface, surface downward solar radiation | 0.25 decimal degrees | global | hourly | 1950 - present | Hersbach et al., 2020 |
| Copernicus Global Land Service Land Cover | Land cover categories | 100m | global | annual | 2015 - two years from present | Buchhorn et al., 2020 |
| Potapov et al. 2020 | Vegetation Canopy Height | 30m | within 52°N and 52°S globally | static | 2019 | Potapov et al., 2020 |
| Simard et al. 2011 | Vegetation Canopy Height | 1000m | global (employed for above 52°N and 52°S) | static | 2005 | Simard et al., 2011 |
| MODIS MOD15A2H | LAI (Leaf Area Index) Stage 2 | 500m | global | 8 days | 2000 - 2020 | Myneni et al., 2015 |
| ISRIC-WISE | Soil bulk density and percent clay, silt, and sand at 0cm, 5cm, 15cm, 30cm, 60cm, and 100cm depths | 0.05 decimal degrees | global | static | spatial data sources from 1970s to 2000s | Batjes et al., 2012 |
| ASTER Global Digital Elevation Map | Elevation (GDEM V3) | 30m | global | static | Launched in 1999 | Fujisada et al., 2005 |

#### WebReferences

Batjes NH. 2012. ISRIC-WISE derived soil properties on a 5 by 5 arc-minutes global grid (ver. 1.2). Wageningen: ISRIC — World Soil Information.

Buchhorn M, Smets B, Bertels L, *et al.* 2020. Copernicus Global Land Service: Land Cover 100m: collection 3: epoch 2019: Globe.

Fujisada H, Bailey GB, Kelly GG, *et al.* 2005. ASTER DEM performance. *IEEE Transactions on Geoscience and Remote Sensing* **43**: 2707–14.

Hersbach, H., Bell, B., Berrisford, P., Hirahara, S., Horányi, A., Muñoz-Sabater, J., Nicolas, J., Peubey, C., Radu, R., Schepers, D., Simmons, A., Soci, C., Abdalla, S., Abellan, X., Balsamo, G., Bechtold, P., Biavati, G., Bidlot, J., Bonavita, M., Chiara, G.D., Dahlgren, P., Dee, D., Diamantakis, M., Dragani, R., Flemming, J., Forbes, R., Fuentes, M., Geer, A., Haimberger, L., Healy, S., Hogan, R.J., Hólm, E., Janisková, M., Keeley, S., Laloyaux, P., Lopez, P., Lupu, C., Radnoti, G., Rosnay, P. de, Rozum, I., Vamborg, F., Villaume, S. & Thépaut, J.-N. (2020) The ERA5 global reanalysis. *Quarterly Journal of the Royal Meteorological Society*, **146**, 1999–2049.

Myneni R, Knyazikhin Y, and Park T. 2015. MOD15A2H MODIS/Terra Leaf Area Index/FPAR 8-Day L4 Global 500m SIN Grid V006. *NASA EOSDIS Land Processes DAAC,*.

Potapov P, Li X, Hernandez-Serna A, *et al.* 2021. Mapping global forest canopy height through integration of GEDI and Landsat data. *Remote Sensing of Environment* **253**: 112165.

Simard M, Pinto N, Fisher JB, and Baccini A. 2011. Mapping forest canopy height globally with spaceborne lidar. *Journal of Geophysical Research: Biogeosciences* **116**.

WebTable 3. Classification differences between macroclimate and microclimate, along with kappa agreement statistics and two-sample Wilcoxon test results between macroclimate and microclimate on the spatial locations of each Köppen-Geiger class (all tests significant at the *p* < 0.05 level). Kappa and Wilcoxon statistics could not be calculated for five classes that were not predicted to occur in the case study regions/latitudinal strips by macroclimate (Dfc, EF), or were not predicted to occur by either macroclimate or microclimate (Dfd, Dsd, Dwd). Amount of agreement according to kappa values is as follows: <0.4 = poor; 0.4 - 0.55 = fair; 0.55 - 0.7 = good; 0.7 - 0.85 = very good; >0.85 = excellent agreement.

| Köppen-Geiger Class | Median Difference | % Cells with Difference | kappa statistic | Wilcoxon Test Statistic | Effect Size | Magnitude of Wilcoxon Test Effect Size |
| --- | --- | --- | --- | --- | --- | --- |
| Af | 0 | 18.6 | 0.86 | 1.10 x 10^12^ | 0.07 | small |
| Am | 1 | 51 | 0.508 | 2.26 x 10^11^ | 0.28 | small |
| As | 8.1 | 86.5 | 0.14 | 4.60 x 10^7^ | 0.33 | moderate |
| Aw | 0 | 33.5 | 0.431 | 5.94 x 10^11^ | 0.25 | small |
| BSh | 1 | 65.6 | 0.299 | 2.07 x 10^12^ | 0.02 | small |
| BSk | 0.1 | 37.8 | 0.227 | 5.83 x 10^11^ | 0.18 | small |
| BWh | 0 | 4 | 0.811 | 1.11 x 10^13^ | 0.04 | small |
| BWk | 1 | 90 | 0.093 | 4.16 x 10^7^ | 0.32 | moderate |
| Cfa | 1 | 60.5 | 0.251 | 4.43 x 10^10^ | 0.47 | moderate |
| Cfb | 0 | 45.1 | 0.543 | 3.25 x 10^10^ | 0.11 | small |
| Cfc | 0 | 16.4 | 0.669 | 3.71 x 10^10^ | 0.15 | small |
| Csa | 0 | 41.5 | 0.555 | 9.68 x 10^11^ | 0.05 | small |
| Csb | 0 | 43.9 | 0.691 | 1.09 x 10^12^ | 0.13 | small |
| Csc | 1 | 61.5 | 0.259 | 1.30 x 10^10^ | 0.04 | small |
| Cwa | 0 | 23.5 | 0.655 | 4.53 x 10^11^ | 0.03 | small |
| Cwb | 0 | 43.8 | 0.508 | 4.28 x 10^10^ | 0.12 | small |
| Cwc | 0 | 33.9 | 0.25 | 3.35 x 10^7^ | 0.06 | small |
| Dfa | 0 | 23.4 | 0.512 | 2.19 x 10^11^ | 0.2 | small |
| Dfb | 0.1 | 50.3 | 0.562 | 2.12 x 10^11^ | 0.31 | moderate |
| Dsa | 0 | 41.4 | 0.322 | 7.38 x 10^10^ | 0.12 | small |
| Dsb | 1 | 80.7 | 0.084 | 1.39 x 10^10^ | 0.13 | small |
| Dsc | 1 | 45.1 | 0.361 | 2.56 x 10^6^ | 0.31 | moderate |
| Dwa | 21.9 | 55 | 0.415 | 4.14 x10^11^ | 0.27 | small |
| Dwb | 0 | 41 | 0.398 | 7.54 x 10^11^ | 0.01 | small |
| Dwc | 0 | 13.8 | 0.565 | 1.25 x 10^10^ | 0.25 | small |
| ET | 18.7 | 45.9 | 0.352 | 1.89 x 10^7^ | 0.19 | small |

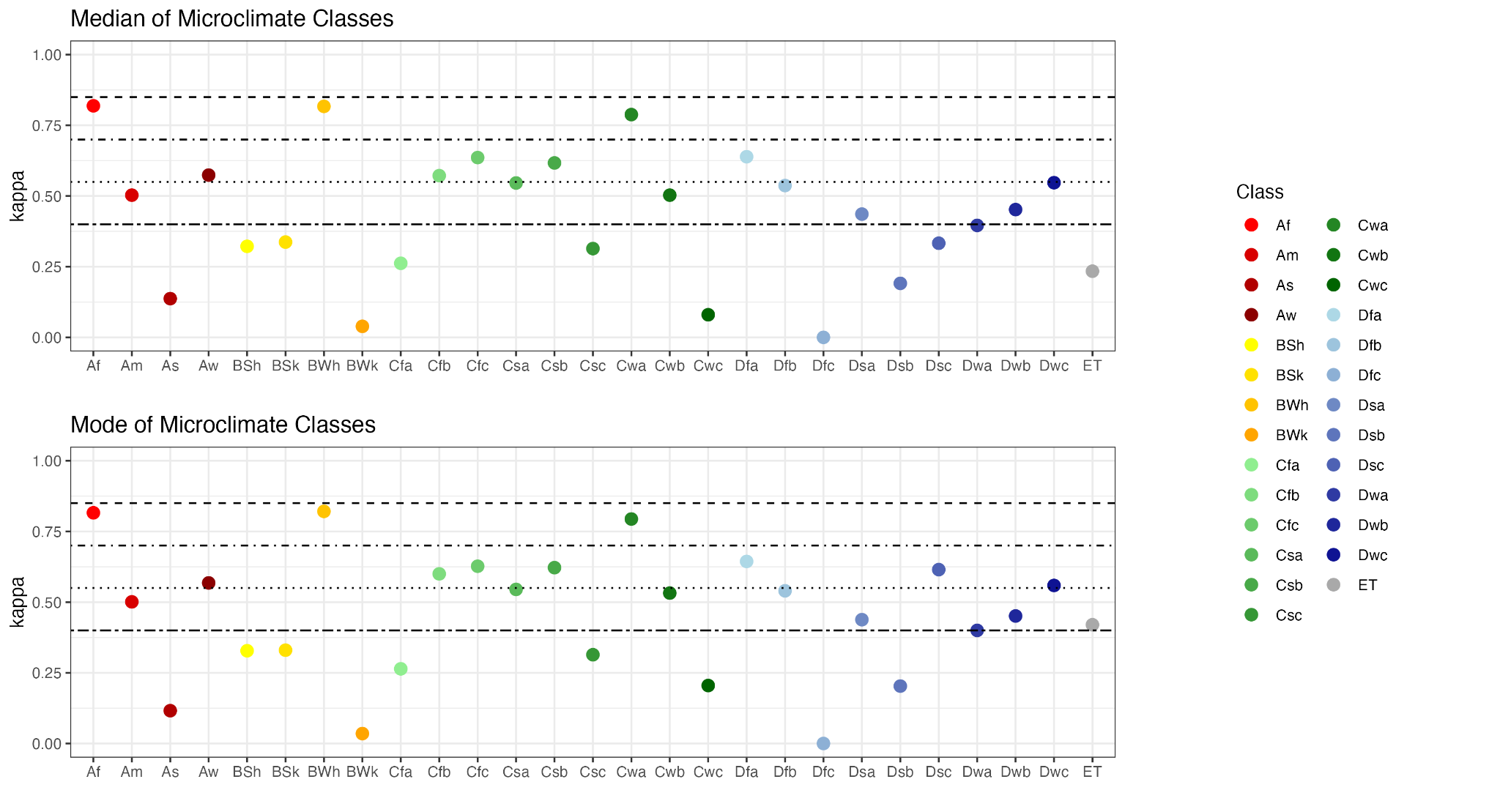

WebFigure 1. Kappa statistics of agreement between macroclimate and microclimate predictions of each Köppen-Geiger class, after microclimate class predictions were aggregated to the median for each macroclimate grid cell (top), and the mode for each macroclimate grid cell (bottom). Amount of agreement according to kappa values is as follows: <0.4 = poor; 0.4 - 0.55 = fair; 0.55 - 0.7 = good; 0.7 - 0.85 = very good; >0.85 = excellent agreement. Similar levels of agreement are indicated here as existed between macroclimate and fine-resolution microclimate (Figure 2, WebTable S3), denoting that the disagreement between macroclimate and microclimate was not because of differences in spatial resolution.

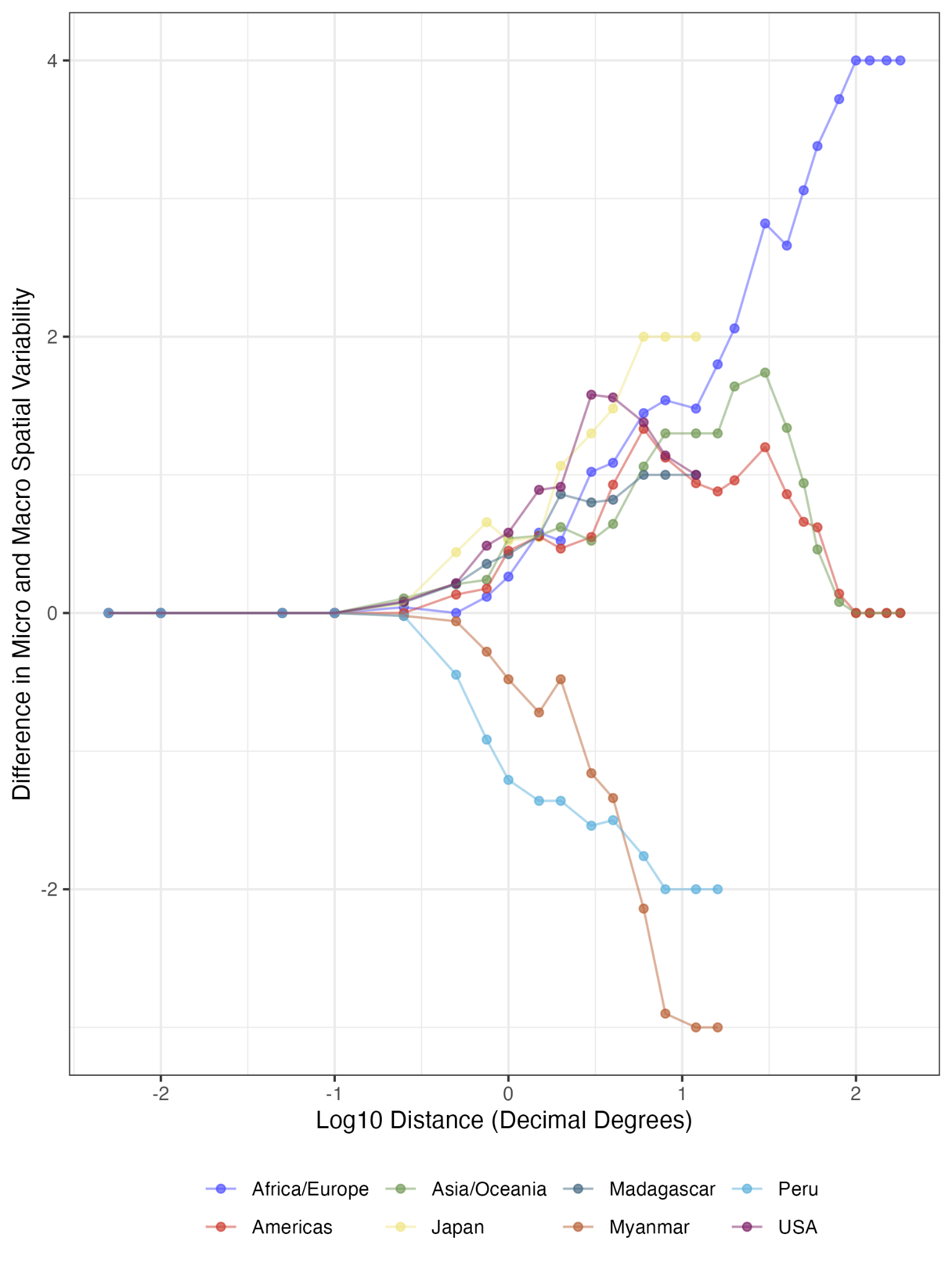

WebFigure 2. Differences in spatial variability (number of classes within a given distance) between macroclimate and microclimate predictions, after microclimate classes were averaged to the median of each macroclimate grid cell (thereby matching the spatial resolution of microclimate). Three out of five study regions, and all three latitudinal strips, demonstrated higher spatial variability in microclimate classes across spatial scales.
